# A lesson from COVID-19 on inaccessibility of web-based information for disabled populations worldwide

**DOI:** 10.1101/2020.08.16.252676

**Authors:** Amiel A. Dror, Eli Layous, Matti Mizrachi, Amani Daoud, Netanel Eisenbach, Nicole G. Morozov, Samer Srouji, Karen B. Avraham, Eyal Sela

## Abstract

Many government websites and mobile content are inaccessible for people with vision, hearing, and cognitive disabilities. The COVID-19 pandemic highlighted these disparities when health authority website information, critical in providing resources for curbing the spread of the virus, remained inaccessible for disabled populations. The Web Content Accessibility Guidelines provide comparatively universally accepted guidelines for website accessibility. We utilized these parameters to examine the number of countries with or without accessible health authority websites. The resulting data indicate a dearth of countries with websites accessible for persons with disabilities. Methods of information dissemination must take into consideration individuals with disabilities, particularly in times of global health crises.

## Introduction

The COVID-19 pandemic is challenging the boundaries of not only social behaviors and cultural institutions, but also the rapid and accurate dissemination of information. The containment of this epidemic has required stringent adherence to interpersonal behavioral modifications, which are often developed and transmitted by national health authorities. However, national health authority websites may lack website accommodations for people with vision, hearing, physical, or cognitive impairments. Minimizing the information gap between able-bodied and disabled populations is imperative for achieving global engagement in containing not only COVID-19, but also future pandemics. We sought to determine what percentage of national health authority websites are fully accessible to people with disabilities according to Web Content Accessibility (WCAG 2.1) guidelines (W3C, 2020a) benchmarks. Our research demonstrates that only a small percentage of government health websites are fully accessible for people with disabilities.

According to the World Health Organization (WHO), an estimated 2.2 billion people suffer from vision impairment or blindness (WHO Blindness and vision impairment, 2019), while 466 million people have a disabling hearing loss (WHO Deafness and hearing loss, 2020). Individuals with temporary or permanent motor or cognitive disabilities also require accessibility modifications for proper interaction with websites. Inconsistent heading level and font size or color contrast of elements in webpages harbor barriers for proper interaction by visually impaired people. Likewise, alternative textual descriptions of visual elements on a page are essential for contextual understanding, in addition to proper interaction with text-to-speech engines. Lack of video content subtitles or transcripts present barriers to the hearing impaired. Compatibility with keyboard navigation, including skip linking in the backend of a website, is crucial to accommodate web navigation for people with motor dysfunction who interact with a single finger or with other motor gestures.

## Materials and methods

The Web Accessibility Initiative (WAI), launched and endorsed by the World Wide Web Consortium (W3C) (W3C, 2020b), established a set of guidelines according to four accessibility principles: whether the website is Perceivable, Operable, Understandable, and Robust. Each WCAG 2.1 principle has a set of testable criteria with a total number of 78 testable success criteria. Each success criteria is assigned to one of three conformance levels: A (lowest), AA (intermediate), and AAA (highest). The adherence to higher levels of conformance has been shown to improve accessibility for both disabled and non-disabled users (Loiacono and Djamasbi, 2013; Ruth-Janneck, 2011; Schmutz *et al*., 2016). A panoply of web accessibility evaluation plug-ins was developed under open-source license for the systematic evaluation of website accessibility against the WCAG 2.1 criteria (W3C, 2020a). A list of available tools is presented by the W3C website without an official recommendation for usage of one tool above another (W3C, 2020c). These automated tools aim to complement the cardinal manual check of a website during the development process and throughout routine website updates to ensure maximal adherence WCAG guidelines (Petrie and Bevan, 2009). A comprehensive comparison between eight widely used accessibility evaluation tools highlights the strengths and weaknesses of each tool and recommends using more than one tool for optimal coverage of success criteria (Frazão and Duarte, 2020). Hence, to test the accessibility of COVID-19 information disseminated through health authority websites, we utilized two independent accessibility evaluation engine including WAVE chrome extension (wave.webaim.org) and Accessibility Insights (accessibilityinsights.io), both of which have been described and utilized in previous literature (Acosta-Vargas *et al*., 2020; Frazão and Duarte, 2020). The WAVE tool analyzes 180 checks according to two conformances level (152 level A; 28 level AA); whereas the Accessibility Insights tool analyzes 64 checks according to three conformances level (55 level A; 7 level AA; and 2 level AAA) (Frazão and Duarte, 2020). It must be noted that the weight of each error (e.g. minor, moderate, critical) is defined by the tool developer and thus may result in different impacts on the overall accessibility rank of the page results.

Due to the rapid growth of COVID-19 information and the frequent updates of health authorities’ websites, which may influence the accessibility score at a given time point, the degree of accessibility of each website was evaluated at three different time points and the presented data refer to the following three consecutive days (5-7 April, 2020). The calculated number of errors of each health authority homepage augments the average number of errors in each test separately (WAVE and Accessibility Insights), with removal of redundant errors represented in both tests.

In addition to accessibility assessments, we tested each website for mobile usability in concordance to Google webmaster developer tools (developers.google.com). In this regard, previous studies have demonstrated that mobile-friendliness of a given website contributes not only to end user usability, but also for website visibility on search engine results (Schubert, 2016).

The list of health authorities’ websites of 189 countries were drawn from The Geneva Foundation for Medical Education and Research (GFMER, 2020) (Supplementary Table 1). Prior to accessibility evaluation, a manual check of each website on the list yielded 174 health authority websites. Websites of 15 countries were excluded due to an inability to load the site on the test server or when the official health authority homepage appeared as a social media page. This was a cross-sectional study concentrating on the accessibility of health authorities’ websites’ homepages (unit of analysis) providing health information and recommended public protective measures against COVID-19.

## Results

Only 4.7% of the countries examined had fully implemented the WAI accessibility guidelines: Italy, the Netherlands, Norway, Japan, Poland, South Korea, the United Kingdom, and the United States (Figure 1). In contrast, sites from the majority of countries continue to have accessibility errors that present significant barriers to people with disabilities around the world. Distribution of reported errors across all 174 tested health authorities’ homepages, according to WCAG conformance levels, reveals that 89% violate Level A criteria, while 11% of countries contain errors that violate higher levels of success criteria (AA and AAA). Inspection of the numbers of errors on all tested pages grouped by WCAG principles indicate that the most impacted principles are robustness (39%) and perceptibility (32%), as compared to operability (19%) and understandability (10%). While both error number, conformance, and principle distribution may be altered according to the selected assessment tools, the data collected signifies the insufficient implementation of WCAG guidelines in the majority of health authority websites, rendering accessibility barriers to millions of people.

**Figure 1.**
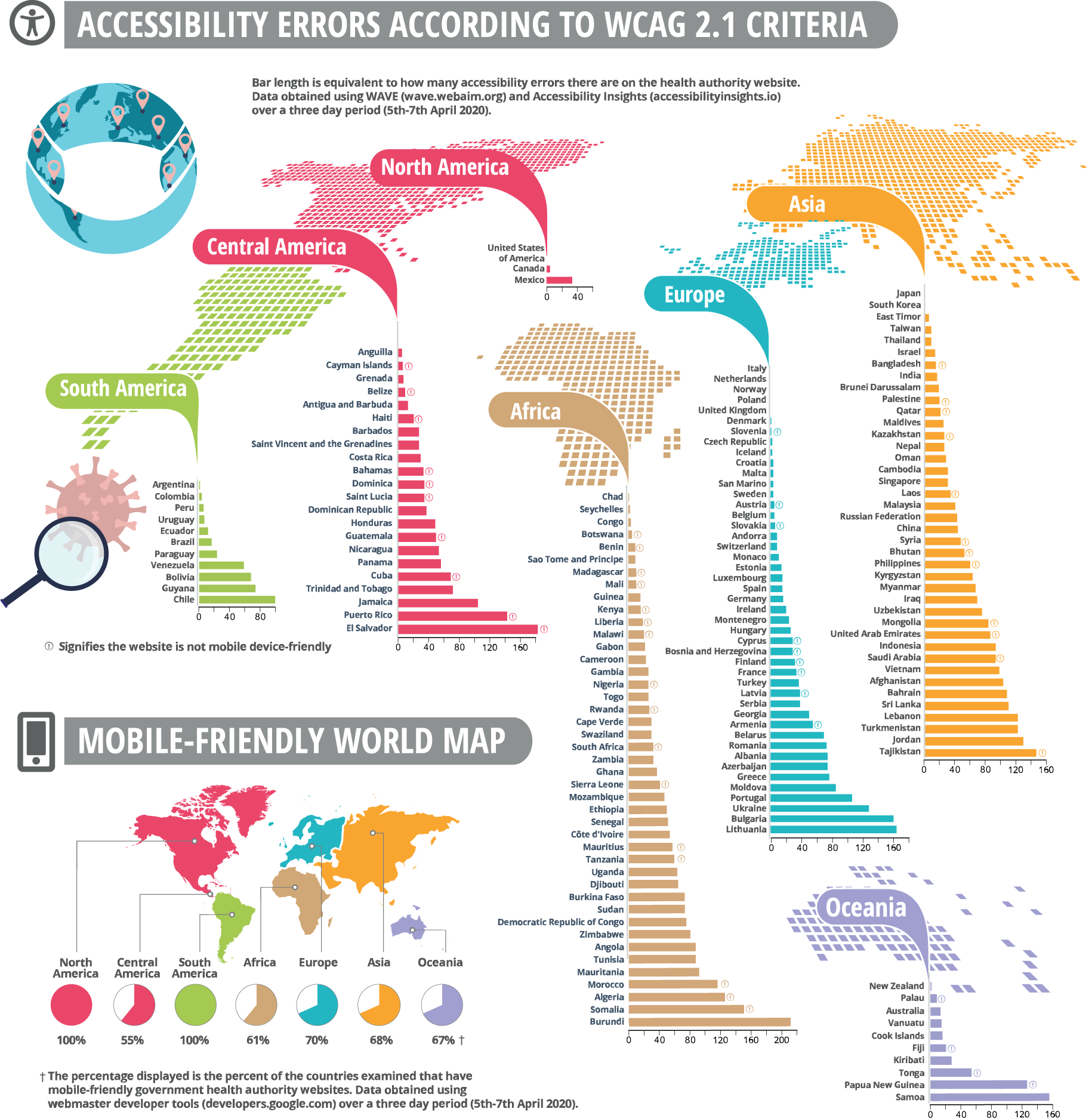
Health authority websites of 174 countries worldwide, demonstrating accessibility errors and mobile friendly maps. A. The calculated number of errors of each health authority website augment the number of error results in each accessibility evaluation tool separately (WAVE and Accessibility Insights), following obliteration of redundant errors that are represented in both tests. B. Mobile computability check according to the Google web developer tool with either pass of fail results. All tests were performed on three consecutive days (5-7 April 2020).

## Discussion

Reducing transmission of SARS-CoV-2 depends on tight adherence of the public to simple but challenging modifications in social and public behavior (Chu *et al*., 2020). Digital media provide numerous platforms to distribute essential information to the public through websites, social media, and instant messaging applications (Jung and Shin, 2020).

Due to the diversity of reporting sources and the harmful consequences of disinformation (Islam *et al*., 2020), governments often encourage the public to check local health authority websites frequently for regular updates. This demand requires the information on official websites to be accessible to as many citizens as possible. Unfortunately, individuals with the greatest need for timely and precise data may have the most difficulty accessing governmental material (West and Miller, 2006). Providing consistently high-quality government productions could also lead to a greater utilization of the Internet by persons with disabilities. Enhancing accessibility to government-sponsored resources could lead not only to immediate population benefits but could also promote the position of disabled people in the digital sphere through increased communication, global engagement, and visibility.

Despite remarkable technological advancements in recent history, for people with visual, hearing, motor and cognitive disabilities, a seemingly simple website interaction can present a daunting challenge. Although internet access is still unavailable to approximately one-third of the world’s population, the needs of all existing users must be accommodated to ensure equal benefits and access to essential health information. The growth and expansion of the Internet must therefore be accompanied by an equal development of sophisticated accessibility technologies, which would expand the usability of the web to individuals with disabilities. Without underestimating the importance of accessibility implementation during normal times, the current COVID-19 pandemic now highlights just how important unhindered access to government websites is during a global health crisis.

